# Trehalose metabolism regulates transcriptional control of muscle development in lepidopteran insects

**DOI:** 10.1101/2025.10.20.683588

**Authors:** Sharada Mohite, Tanaji Devkate, Prashant Kalaskar, Prashant Singh, Abhishek Subramanian, Rakesh Shamsunder Joshi

**Affiliations:** Biochemical Sciences Division, CSIR-National Chemical Laboratory, Dr. Homi Bhabha Road, Pune, 411008, Maharashtra, India; Academy of Scientific and Innovative Research (AcSIR), Ghaziabad 201002, Uttar Pradesh, India; Department of Biotechnology, Indian Institute of Technology Hyderabad, Kandi, Sangareddy, Telangana, 502284, India

**Keywords:** *Helicoverpa armigera*, muscle development, *Spodoptera frugiperda*, transcription factors, trehalose metabolism

## Abstract

The connection between trehalose metabolism and insect muscle development has long been unclear. Here we demonstrate that trehalose-derived glucose flux is essential for sustaining glycolysis and maintaining cellular energetics during metamorphosis in *Helicoverpa armigera*. Disruption of trehalose synthesis through silencing of trehalose 6-phosphate synthase/phosphatase (*TPS/TPP*) or paramyosin (*Prm*) produced defective eclosion and alike fragmented muscle fibers. Metabolic profiling revealed broad depletion of glycolytic intermediates and cofactors consistent with elevated AMP and activation of energy-stress signalling. Transcriptome analysis also showed downregulation of the transcription factor *E2F/Dp*, along with reduced expression of *cyclin/CDK* components, indicative of cell cycle checkpoint dysregulation. *E2F/Dp* repression resulted in lower expression of *Mef2, Prm, MHC,* and other sarcomeric components, causing defective muscle development and weakened muscle architecture. Gene set enrichment analysis, gene regulatory network and promotor binding analysis confirmed E2F binding motifs within promoters of trehalose metabolism and myogenic genes, suggesting a potential metabolic–transcriptional connection. Furthermore, dietary trehalose supplementation partially rescued metabolic depletion and restored expression of *E2F* targets in *TPS/TPP*- and *E2F/Dp*-silenced insects. Collectively, our findings establish trehalose metabolism as a metabolic regulator that couples energy homeostasis to cell-cycle transcriptional control and muscle development.

## 1. Introduction

Insects are the most diverse animal group on earth, and their ecological success depends on tightly regulated developmental programs, particularly the formation of contractile muscles that underlie locomotion, feeding, flight, mating, and successful eclosion (Bretscher & O’Connor, 2020; Denlinger & Zdárek, 1994; Iwamoto, 2011; Jankielsohn, 2018). Muscle development is metabolically demanding, requiring coordinated cell proliferation, myofibril assembly, and structural maintenance (Bretscher & O’Connor, 2020; Luis & Schnorrer, 2021). Because these processes depend on high ATP turnover, maintaining energy homeostasis is a prerequisite for robust muscle development (Bretscher & O’Connor, 2020).

Carbohydrates serve as the primary fuel for insect development. In *Drosophila melanogaster*, impaired glycolysis or glycogen synthesis compromises ATP production and leads to thin or degenerated muscle fibers (Bawa et al., 2020; Tixier et al., 2013; Yamada et al., 2018). In addition to glucose and glycogen, trehalose, the primary sugar in insects, is crucial for muscle function (Sacktor & Wormser-Shavit, 1966). Trehalose is synthesized in the fat body by trehalose 6-phosphate synthase/phosphatase (TPS/TPP) and hydrolyzed to glucose by trehalase (Treh) at target tissues (Shukla et al., 2015; Tang et al., 2012). Trehalose not only fuels muscle contractility but is indispensable for eclosion, when coordinated contractions enable adults to emerge from the pupal case (Bothe & Galler, 1996; Kammer & Kinnamon, 1977). Indeed, loss of *Tps1* in *D. melanogaster* or inhibition of TPP in *Helicoverpa armigera* causes pupal lethality due to failed emergence (Matsuda et al., 2015; Tellis et al., 2024). Despite these observations, the mechanistic link between trehalose metabolism and muscle development remains unresolved.

Emerging evidence suggests that metabolic state directly influences transcriptional networks. Insects, like mammals, use nutrient-derived signals to regulate transcription factors such as carbohydrate response element binding protein (ChREBP) and max-like protein X (Mlx) that connect glucose availability to gene expression (Bravo-Ruiz et al., 2021). Whether trehalose metabolism similarly couples cellular energetics to transcriptional control of muscle development has not been established.

To investigate the relationship between trehalose anabolism and muscle development in *H. armigera* (Ha), we employed a comprehensive and integrative methodology. By performing transcriptional and small-molecule inhibitory perturbations in TPS/TPP, we observe a multitude of molecular and phenotypic changes. Our results establish the involvement of TPS/TPP in trehalose metabolism, altering global myogenic gene expression, development of muscle filaments and their growth. Prediction of underlying gene regulatory network and gene set enrichment analysis from the *H. armigera* transcriptome, as well as gene expression analysis, suggests putative TFs that may regulate the expression of myogenic genes in response to organismal trehalose levels. Partial restoration of TFs, downstream gene expression and trehalose biosynthesis by exogenous trehalose in *HaTPS/TPP* and TF-silenced insects further confirms the hypothesis of the regulatory role of trehalose metabolism in insect muscle development. Our study thus establishes trehalose metabolism as a central regulator that couples energy homeostasis to cell-cycle transcriptional control and muscle development.

## 2. Results

### 2.1 Inhibition of TPP dysregulates trehalose metabolism and affects myogenic genes, resulting in late pupal mortality

Disruption of trehalose synthesis by N-(phenylthio) phthalimide (NPP) feeding harms the trehalose metabolism and overall growth and development of lepidopteran insects (Tellis et al., 2024). We observed that NPP feeding disrupted trehalose synthesis, resulting in delayed pupation and failed eclosion **(Figure 1A)**. To investigate these developmental defects, we examined the expression levels of genes involved in trehalose metabolism and muscle development upon TPP inhibition. Muscle development rely on the robust expression of structural genes, we studied several myogenic genes including *myosin heavy chain* 95F (*MHC95F*), *actin88F* (*Act88F*), *myosin regulatory light chain* 2 (*MLC2*), *flightin (Fln)*, *bent*, *paramyosin (Prm)*, *tropomyosin 2 (Tm2)*, *troponin C (TpnC)*, *myocyte enhancer factor 2* (*Mef2*) and *alpha-actinin* (Hooper & Thuma, 2005; Zappia & Frolov, 2016).

**Figure 1.**
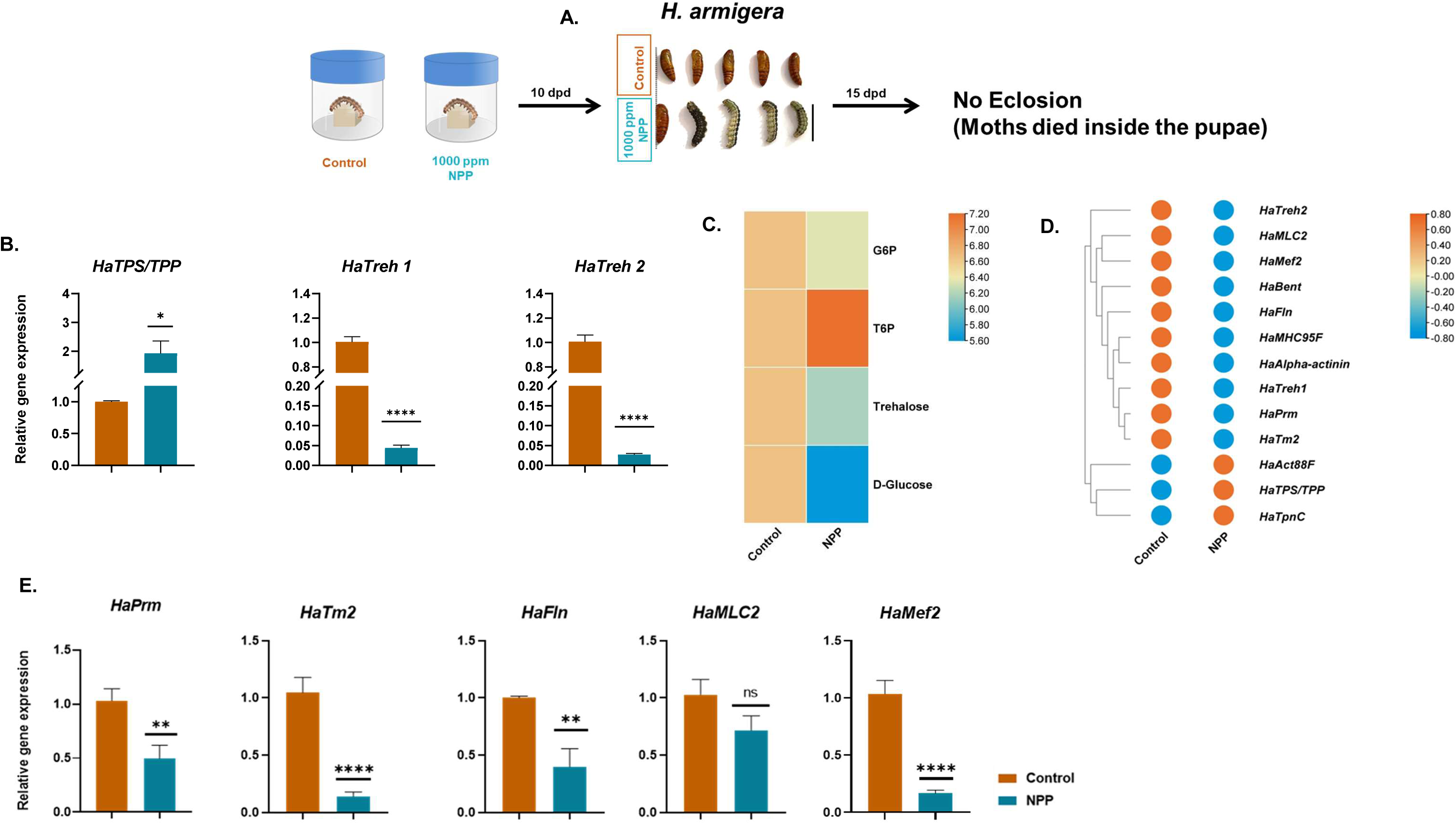
Impact of NPP feeding on the growth, development, and gene expression of *Helicoverpa armigera* and *Spodoptera frugiperda*. (A) Schematic representation of NPP feeding assay with *H. armigera* larvae carried out for 10^th^ days (n = 100, second-instar larvae, dpd= days post diet). Representation of insects on the 10^th^ day and 15^th^ day of NPP feeding bioassay, indicating delayed growth, pupation, reduced size, and no eclosion. (B) Gene expression analysis of trehalose metabolism genes using qRT-PCR. (C) Relative quantification of metabolites formed in trehalose metabolism. (D) Differential expression analysis of genes involved in trehalose metabolism and muscle development using NPP-fed insect transcriptomics data. Genes are clustered using hierarchical clustering. (E) Gene expression analysis of selected myogenic genes in control and treatment using qRT-PCR. Asterisks indicate the significance level between control and treatment. Student’s unpaired t-test was used for calculating significance. Data are represented as mean ±SEM; (*p<0.05; **p value <0.01; *** p value <0.001; ****p value <0.0001; ns= non-significant).

Comparison of gene expression in control vs. NPP-fed insects indicated a significant upregulation of *HaTPS/TPP* (∼2-fold increase) while a significant downregulation of *HaTreh 1* and *HaTreh 2* (∼9-fold decrease) **(Figure 1B)**. Furthermore, NPP feeding leads to reduced levels of energy sources, trehalose, and glucose **(Figure 1C)**, indicating dysregulation of energy metabolism upon TPP inhibition. Differential gene expression analysis on the *de novo* bulk RNA sequencing data obtained from control vs. NPP-fed insects analysis showed that most of the myogenic genes are deregulated **(Figure 1D)**. Validation by real-time PCR (qRT-PCR) indicated that the myogenic genes such as *HaPrm, HaTm2, HaFln, and HaMef2* showed significant downregulation while *HaMLC2* seemed inconsistent albeit decreasing trend **(Figure 1E)**, upon NPP feeding. These findings collectively indicate that disruption of trehalose synthesis might indeed affect the growth, development, and maintenance of muscles in *H. armigera*.

### 2.2 Silencing of *TPS/TPP* resulted in disturbed trehalose metabolism and muscle development

To eliminate any potential influence of inhibitor feeding on eclosion, we employed a dsRNA-based approach to silence the *TPS/TPP* gene in *H. armigera* and *Spodoptera frugiperda* (Sf) and examined their effects on muscle development **(Figure 2-figure supplement 1)**. *HaTPS/TPP* silencing resulted in a significant decrease in *HaTPS/TPP* expression (∼50%) **(Figure 2A),** and a reduction (∼50%) in the expression of *HaTreh 1* and *2* **(Figure 2B)**. Furthermore, silencing led to ∼70 and 60% decrease in TPP and Treh activities, respectively **(Figure 2A-B)**. This reduction caused lower levels of trehalose and glucose **(Figure 2C)**, indicating a disruption in trehalose metabolism.

**Figure 2.**
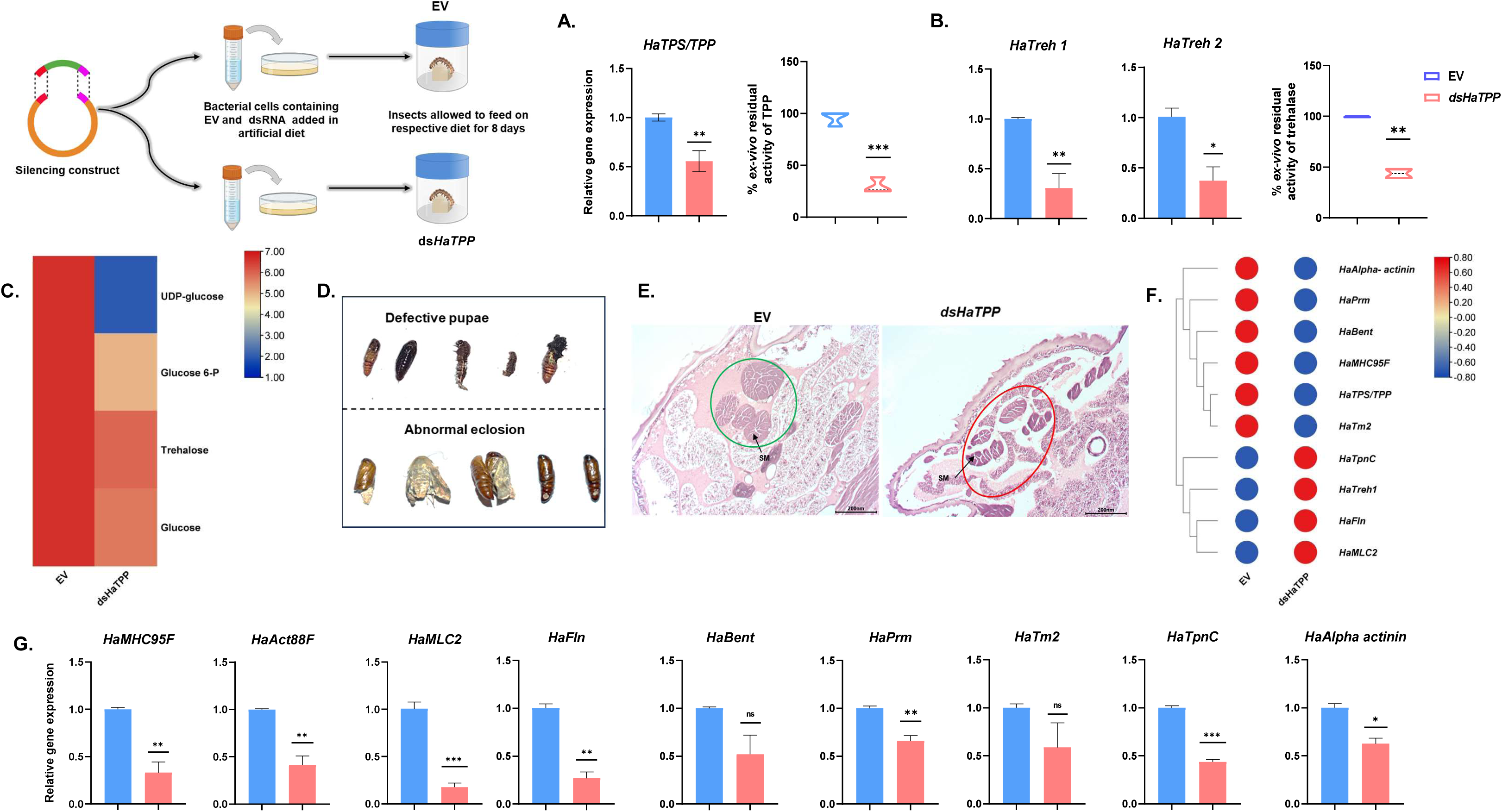
dsRNA mediated silencing of *H. armiger*a *HaTPS/TPP.* Schematic representation of dsRNA feeding assay carried out for 8^th^ days (n = 60, second-instar larvae). (A) Silencing confirmation of *HaTPS/TPP* using qRT-PCR and percent ex-vivo residual activity of *Ha*TPP. (B) Gene expression of *HaTreh1* and *HaTreh2* using qRT-PCR and percent ex-vivo residual activity of *Ha*Treh. (C) Relative quantification of metabolite level of trehalose metabolism in *HaTPS/TPP*-silenced insects of *H. armigera*. (D) Phenotypic abnormalities associated with pupae and moths were observed upon *dsHaTPP* feeding. (E) Histological analysis of longitudinal sections (∼8um) fifth instar larvae of *H. armigera* upon feeding of EV and *dsHaTPP*. Scale bar=200nm. (F) Heatmap representing the expression pattern of genes involved in trehalose metabolism and muscle development upon *HaTPS/TPP* silencing. (G) Gene expression analysis of selected myogenic genes upon *dsHaTPP* feeding by qRT-PCR. Asterisks indicate the significance level between control and treatment. Student’s unpaired t-test was used for calculating significance. Data are represented as mean ±SEM; (*p<0.05; **p value <0.01; *** p value <0.001; ****p value <0.0001; ns= non-significant).

Despite this metabolic disruption, larvae fed a diet containing *dsHaTPP* exhibited no significant changes in body mass or size compared to control. However, ∼40% mortality was observed in the *HaTPS/TPP*-silenced larvae **(Figure 2-figure supplement 2A)**. Interestingly, silencing also led to defective pupae and abnormal eclosion, highlighting its detrimental effect on larval and pupal development **(Figure 2D)**. Given these abnormalities in eclosion, we performed histological analysis to assess muscle structure. *HaTPS/TPP*-silenced larvae displayed irregular and fragmented skeletal muscles (SM) compared to EV-fed larvae **(Figure 2E)**, suggesting impaired muscle development.

To further validate this finding, we conducted transcriptomic analysis following *dsHaTPP* feeding. Differential gene expression analysis revealed that many myogenic genes, such as *HaAlpha-actinin*, *HaPrm, HaBent, HaMHC95F*, and *HaTm2*, followed the decreasing expression pattern of *HaTPS/TPP*. In contrast, genes like *HaTpnC, HaFln*, and *HaMLC2* mirrored the expression pattern of *HaTreh1* **(Figure 2F)**. Further validation through qRT-PCR demonstrated that reduced expression of myogenic genes, including *HaMHC95F, Haact88F, HaMLC2, HaFln, HaPrm, HaTnC,* and *HaAlpha-actinin* in *HaTPS/TPP* silenced insects **(Figure 2G)**. Overall, these findings suggest that the inhibition of trehalose synthesis negatively impacts *H. armigera* muscle development.

Similarly, in *S. frugiperda* (Sf), ds*SfTPS/TPP* feeding resulted in a significant reduction in its expression (∼40%) and activity (∼30%) **(Figure 2-figure Supplement 3A)**. Furthermore, *SfTreh* showed a notable decrease in expression (∼65%) and activity (∼20%) **(Figure 2-figure supplement 3B)**. However, body weight or size were observed to be unchanged in the ds*SfTPP*-fed larvae **(Figure 2-figure supplement 3C)**. Based on these observations, we hypothesized that the reduction in trehalose and glucose levels might also impact *S. frugiperda* muscle development. *SfPrm* and *SfTm2* showed a significant decrease in expression (∼1 to 1.5-fold) upon silencing of *SfTPS/TPP* **(Figure 2-figure supplement 3D)**. In conclusion, our findings demonstrate that the *H. armigera* and *S. frugiperda TPS/TPP* silencing disrupts trehalose metabolism, negatively affects larval-pupal muscle development, and leads to unsuccessful eclosion, suggesting a potential connection between trehalose synthesis and muscle development.

### 2.3 *HaTPS/TPP* silencing affect muscle development that mimics with muscle phenotype on *HaPrm* silencing

To confirm the changes in muscle structure and abnormal phenotypes observed after inhibiting trehalose synthesis, we silenced the *Prm*, an invertebrate-specific muscle gene, in *H. armigera*. Feeding larvae with the *dsHaPrm* resulted in ∼40% silencing of *HaPrm* **(Figure 3A)**. However, body mass or size of larvae fed on a diet containing *dsHaPrm* were unaltered. The mortality rate of *HaPrm*-silenced larvae was ∼35% **(Figure 3-figure supplement 2B)**. However, *HaPrm* silencing led to morphological defects in larvae, pupae, and abnormal wings in adult moths, suggesting disrupted metamorphosis **(Figure 3B)**. Histological analysis revealed fragmented and irregularly shaped skeletal muscles in *HaPrm*-silenced insects **(Figure 3C)**.

**Figure 3.**
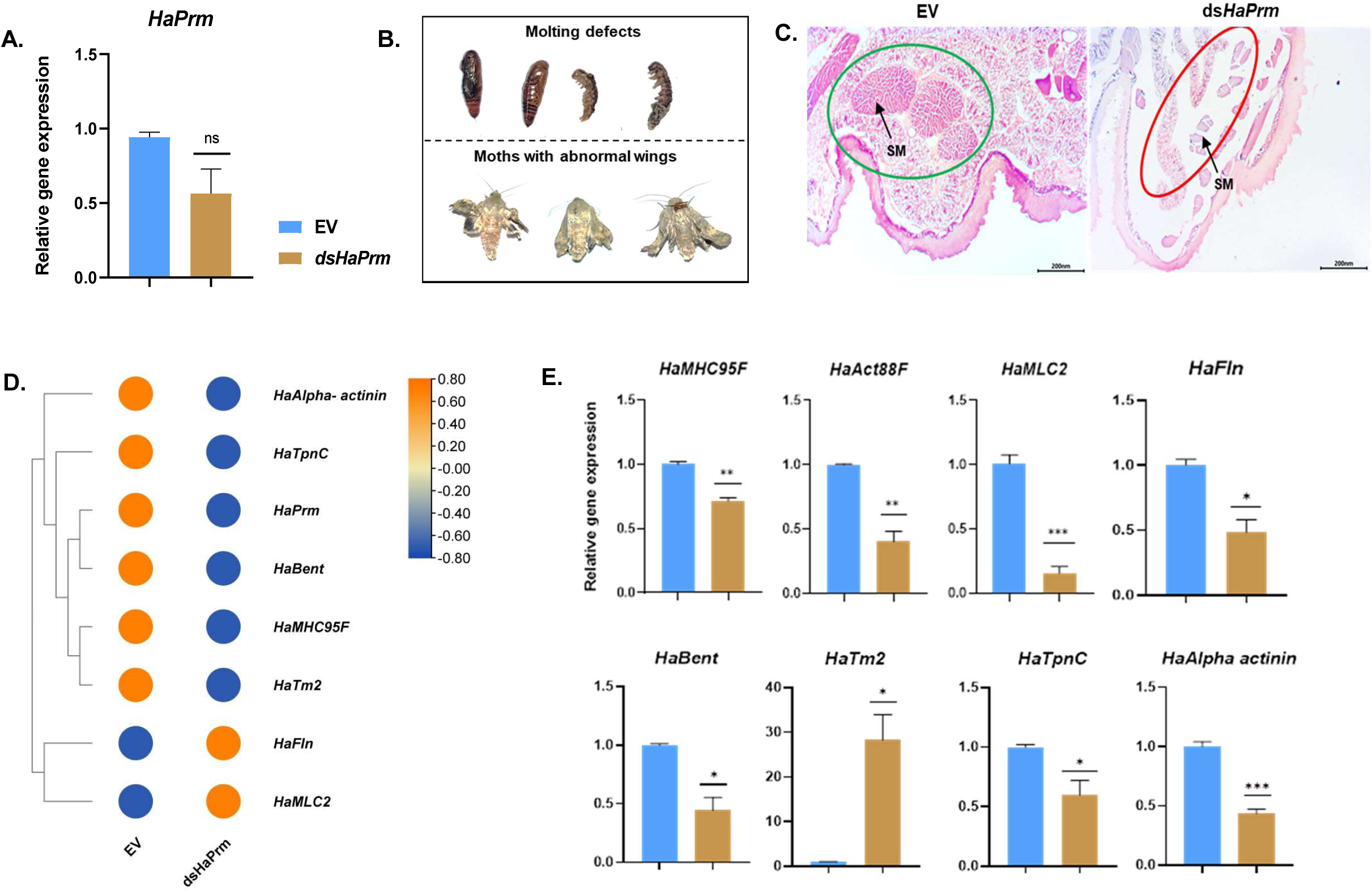
dsRNA mediated silencing of *H. armiger*a *Prm* (n=60, Second instar larvae). (A) Silencing confirmation of *HaPrm* using qRT-PCR. (B) Phenotypic abnormalities associated with pupae and moths were observed upon *dsHaPrm* feeding. (C) Histological analysis of longitudinal sections (∼8um) fifth instar larvae of *H. armigera* upon feeding of EV and *dsHaPrm*. Scale bar=200nm. (D) Heatmap represents the gene expression pattern involved in muscle development upon *HaPrm* silencing. (E) Gene expression analysis of myogenic genes upon *HaPrm* silencing using qRT-PCR. Asterisks indicate the significance level between control and treatment. Student’s unpaired t-test was used for calculating significance. Data are represented as mean ±SEM; (*p<0.05; **p value <0.01; *** p value <0.001; ****p value <0.0001; ns=non-significant).

To investigate further, we examined transcriptional changes through *de novo* bulk RNA sequencing of *HaPrm*-silenced insects and performed differential gene expression profiling of various myogenic genes. Most of the myogenic genes were downregulated following *HaPrm* silencing **(Figure 3D)**. This was confirmed by qRT-PCR, which showed a significant reduction in gene expression **(Figure 3E)**. These findings indicate that muscle defect upon silencing *HaPrm* is similar to defects in *TPS/TPP* silenced *H. armigera*.

### 2.4 Trehalose metabolism disruption induces altered glycolytic pathways, energy depletion and cell cycle deregulation

*HaTPS/TPP* silencing leads to a reduction in trehalose and glucose levels, prompting further analysis of metabolites and gene expression involved in glycolytic and related metabolic pathways. Following *HaTPS/TPP* silencing, we observed a significant decrease in the levels of key intermediates in the glycolytic and hexosamine pathways, including fructose 6-phosphate, 3-phospho-D-glycerate, phosphoenolpyruvate, pyruvate, glucosamine, and glucosamine 6-phosphate **(Figure 4A)**. Additionally, there was a notable reduction in energy molecules such as adenosine diphosphate (ADP), nicotinamide adenine dinucleotide (NAD), nicotinamide adenine dinucleotide (NADH), and nicotinamide mononucleotide (NMN), all of which are essential for efficient energy production and utilization **(Figure 4A)**. Furthermore, the expression of key glycolytic genes, *HaEnolase* (*HaENO1*) and *HaPyruvate Kinase* (*HaPK*), was significantly downregulated. In contrast, *HaPhosphoglycerate Kinase* (*HaPgk*) was significantly upregulated **(Figure 4B)**, suggesting a disruption in overall energy metabolism. Next, to investigate the effect of low levels of trehalose and glucose on the cell cycle, we conducted differential gene expression analysis of cell-cycle and energy-sensing genes upon *HaTPS/TPP* silencing. Given that muscle development is tightly regulated by the precise coordination of the cell cycle and metabolism (Miyazawa & Aulehla, 2018), we observed that silencing *HaTPS/TPP* significantly reduced the expression of several key cell cycle regulators, including *S-phase cyclin A-associated protein in the endoplasmic reticulum* (*SCAPER*), *cyclin-dependent kinase 1*, *cyclin-A2-like*, and *G2/mitotic-specific cyclin-B*, indicating a cell cycle deregulation (Britton & Edgar, 1998) **(Figure 4C)**. Additionally, we found upregulation of the energy-sensing gene *arginine kinase*, which may help to maintain ATP levels under low energy conditions (Wang et al., 2009) **(Figure 4C)**. Further, we performed expression analyses of transcription factors (TFs) known to be involved in muscle development in response to glucose metabolism. Differential gene expression and qRT-PCR analyses revealed a significant reduction in the expression of *Myocyte enhancer factor 2* (*HaMef2*), *Max-like protein X (HaMlx*), *Mlx interacting protein* (*HaMlxIP*), and *Insulin growth factor 2 (HaIGF2*) following the silencing of *HaTPS/TPP* and *HaPrm* **(Figures 4D)**. However, no changes were observed in the expression of *Forkhead box O* (*HaFoxo*) under the same conditions. Similarly, in *S. frugiperda,* silencing *SfTPS/TPP* led to a notable decrease in the expression of glucose responsive TF, *SfMlx* and *SfMlxIP* **(Figure 4-figure supplement 3E)**. Collectively, these results suggest that trehalose derived glucose and overall cell energetics are crucial for the normal progression of the cell cycle during muscle development.

**Figure 4.**
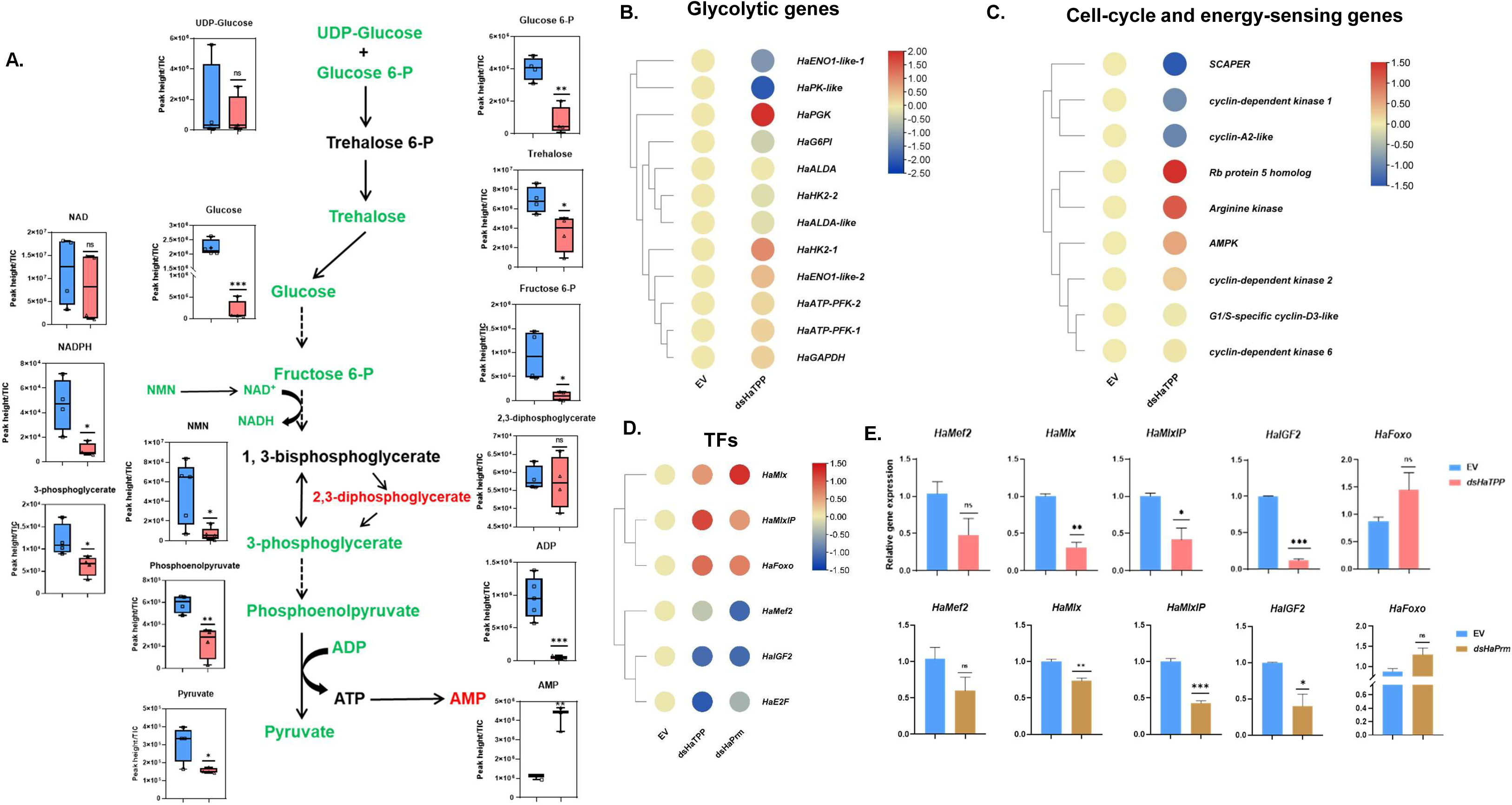
Inhibition of trehalose metabolism disrupts energy homeostasis and affects the regulation of the cell cycle and transcription factor. (A) Schematic of the flow of trehalose and glycolysis toward pyruvate. Metabolites that are significantly reduced upon *HaTPS/TPP* silencing compared to controls are shown in green and increased metabolite shown in red. (B) Heatmap represents the differential gene expression analysis of glycolytic genes upon *HaTPS/TPP* silencing. (C) Heatmap represents the differential gene expression analysis of cell cycle genes and energy-sensing genes upon *HaTPS/TPP* silencing. (D) Differential gene expression TFs upon silencing of *HaTPS/TPP* and *HaPrm*. (E) Validation of gene expression level of TFs by real-time PCR upon *HaTPS/TPP* and *HaPrm* silencing. Asterisks indicate the significance level between control and treatment. Student’s unpaired t-test was used for calculating significance. Data are represented as mean ±SEM; (*p<0.05; **p value <0.01; *** p value <0.001; ****p value <0.0001; ns=non-significant).

### 2.5 Pathway enrichment analysis reveals perturbations in key biological processes upon *TPS/TPP* silencing

Gene Set Enrichment Analysis (GSEA) was performed using the differentially expressed genes and compared with KEGG gene sets to identify significantly enriched biological processes upon TPS/TPP silencing, providing functional context to the expression changes and enabling pathway-level interpretation of regulatory perturbations (Hung et al., 2012; Wixon & Kell, 2000) **(Figure 5A)**. This pathway-focused approach is crucial for understanding how transcriptional changes may translate to phenotypic outcomes. The analysis revealed 11 significantly enriched pathways **(Figure 5B-Supplementary Table S2)**. The normalized enrichment scores (NES) range from −1.8 to 1.7, with positive NES values indicating upregulated processes and negative values representing downregulated processes. Developmental processes like proteasome function, motor protein activity and energy or protein biomass-related metabolic pathways like valine, leucine, and isoleucine metabolism, citric acid cycle, arginine biosynthesis and pyruvate metabolism are transcriptionally upregulated upon TPS/TPP silencing. Similarly, developmental processes like neuroactive ligand-receptor interactions, longevity regulating pathways and metabolic pathways like retinol metabolism, pentose and glucuronate interconversions and metabolism of xenobiotics by cytochrome P450 are transcriptionally downregulated upon TPS/TPP perturbations. Overall, GSEA demonstrates major transcriptional perturbations in metabolic pathways and hypothesized TPS/TPP-induced developmental perturbations **(Figure 5B)**.

**Figure 5.**
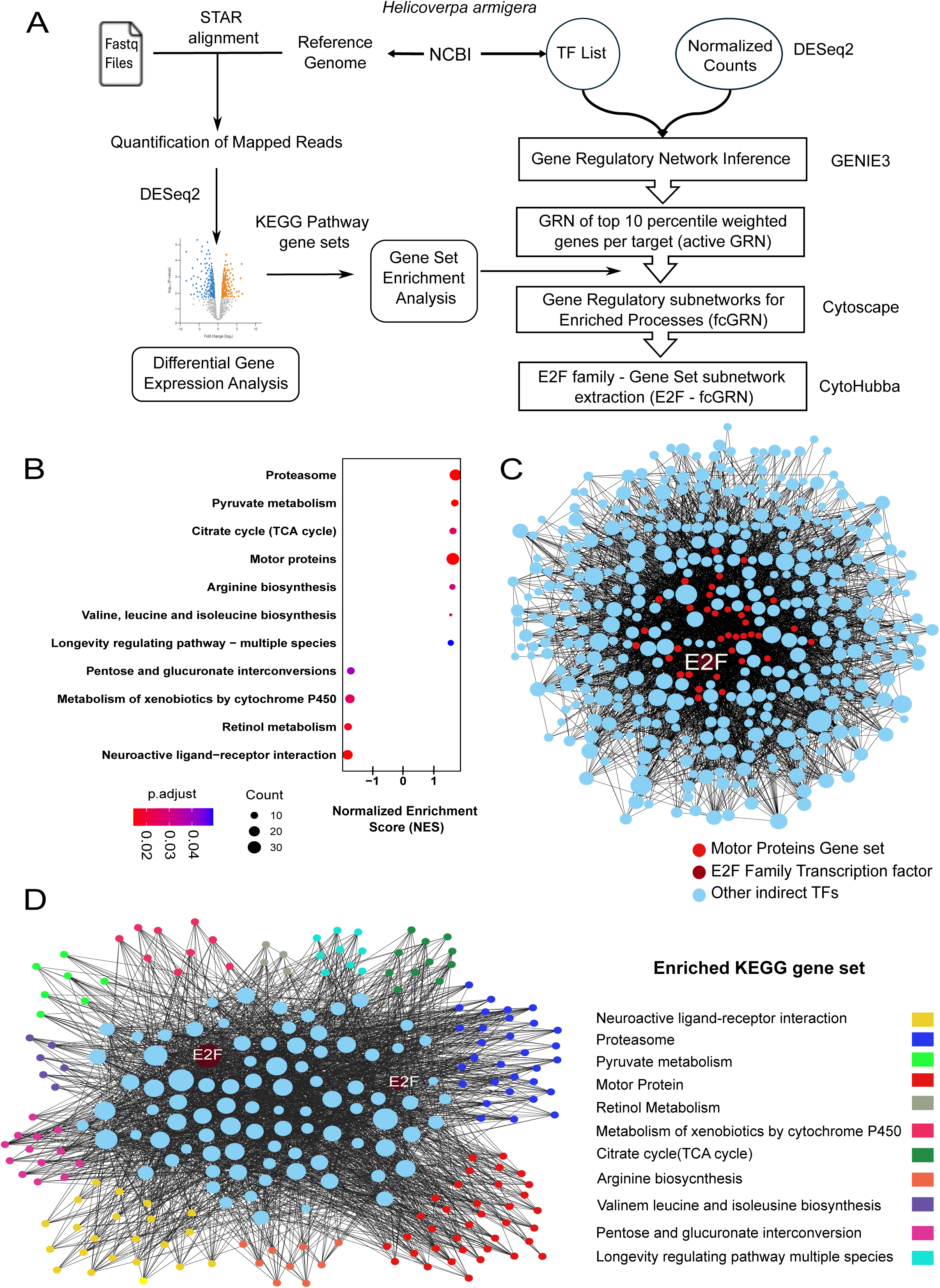
Functional gene regulatory network inference of the TPS/TPP silencing in *H. armigera* (A) Computational pipeline for RNA-seq data processing and network analysis. (B) KEGG pathways enriched upon TPS/TPP silencing, as identified by GSEA. Color scale is dependent upon adjusted P-value. The size of the circles is dependent on the number of enriched genes within each pathway. The X-axis shows the normalized enrichment scores (NES) of the different processes. (C) Gene regulatory subnetwork underlying the enriched motor protein KEGG gene set (motor protein - fcGRN). The network displays transcription factors with an E2F family transcription factor prominently featured at the centre, and their regulatory targets. Node size reflects the number of outgoing connections within the network such that TF with more number of connections will be highlighted. E2F family transcription factor is labelled in maroon, motor proteins are labelled in red and other proteins have a light-blue colour. (D) E2F family – enriched Gene set regulatory subnetwork. The source nodes are E2Fs and the target nodes are the genes belonging to 11 enriched gene sets. The network shows E2F transcription factor at the centre with regulatory connections to genes involved in various KEGG pathways. Nodes are color-coded according to their associated biological pathways as indicated in the legend.

### 2.6 Gene regulatory network analysis predicts E2F family transcription factors to be involved in the regulation of the perturbed gene sets

400 manually curated TFs associated with 571 NCBI gene entries were used to infer active and fcGRNs. The active GRN obtained from our GRN inference methodology was composed of 93,67,712 TF-target interactions. A subnetwork extraction methodology yielded 11 fcGRNs corresponding to the GSEA-enriched KEGG gene sets. We found that the fcGRN corresponding to the motor protein gene set which contains various motor proteins like myosin, kinesin, dynein, tubulin, etc. showed links to many TFs directly or indirectly **(Figure 5C)**. Interestingly, motor proteins are previously implicated in insect muscle function (Iwamoto, 2011). Performing a simple outdegree centrality calculation (outgoing connectivity of nodes with other nodes) within this motor protein fcGRN indicated that an E2F family transcription factor occurs to be a central node (4^th^ rank in the motor protein GRN) involved in this function. Extracting the E2F family TFs – Enriched Gene Set subnetworks we observe E2F family TFs to be connected with the 11 enriched processes (**Figure 5D**). The mRNAs of these implicated E2F family TFs (XM_021328198.3, XM_049842870.2, XM_064034818.1, XM_064034819.1) were obtained from NCBI Gene and shortlisted for experimental verifications.

### 2.7 E2F/Dp regulates the trehalose synthesis and muscle development

Previous studies suggest that E2F1 directly activates the expression of genes involved in glycolysis and later stages of myogenesis during muscle development. Additionally, *E2F1-*deficient *D. melanogaster* larvae exhibited mortality at the pupal stage, suggesting a crucial role for E2F1 in adult skeletal muscle function and overall viability (Zappia et al., 2023a; Zappia & Frolov, 2016). The serendipitous discovery of E2F within the GRN further urged us to explore the putative role of E2F in regulating the expression of myogenic genes in response to fluctuating trehalose or glucose levels. To confirm this, we performed an expression analysis of *E2F* and its dimerization partner *Dp* in *H. armigera* following NPP, *dsHaTPP*, and ds*HaPrm* treatments. Both *E2F* and *Dp* showed significantly reduced expression following silencing of *HaTPS/TPP* and *HaPrm* **(Figures 6A-B)**. However, only E2F expression was notably reduced following NPP feeding, while *Dp* expression showed no significant change **(Figure 6C)**. Given the role of E2F in muscle development and metabolic regulation (Zappia & Frolov, 2016), we predicted the putative binding sites in genes involved in metabolism and muscle development. The transcription factor-binding site analysis of the 5’ UTR regions of *HaTPS/TPP*, *HaTreh*, *HaPgk*, *HaPrm, HaAct88F*, *HaFln,* and *HaTm1* showed the presence of *HaE2F* binding sites **(Figure 6-figure supplement 4A)**. In Electrophoretic Mobility Shift Assay (EMSA), upon incubation with the *Ha*E2F protein, the *HaTPS/TPP* promoter produced progressively retarded bands as the protein concentration increased, indicating dose-dependent formation of *Ha*E2F-*TPS*/*TPP* promoter complexes **(Figure 6D)**. Collectively, these results support a direct regulatory role for E2F in controlling trehalose synthesis genes and related expression programs, including myogenic genes.

**Figure 6.**
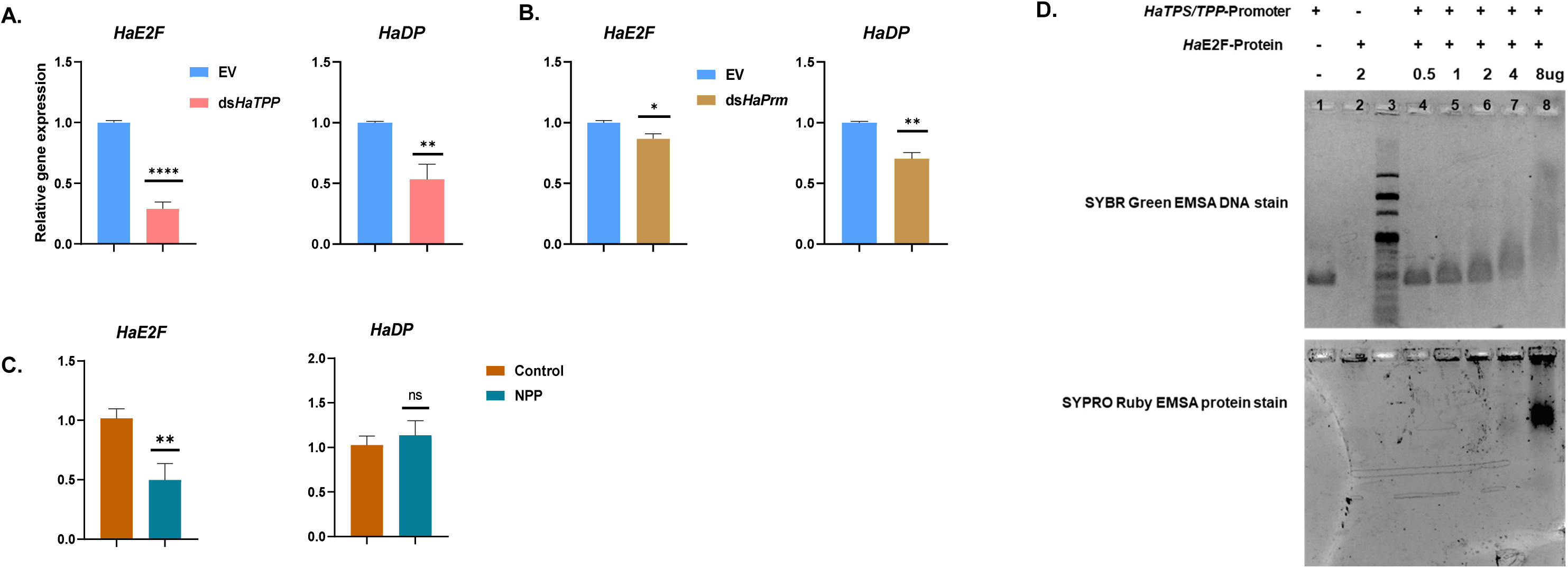
Gene expression analysis and EMSA. (A) Gene expression analysis of *E2F* and *Dp* upon *HaTPS/TPP* silencing. (B) Gene expression analysis of *E2F* and *Dp* upon *HaPrm* silencing. (C) Gene expression analysis of *E2F* and *Dp* upon feeding of 1000 ppm NPP to *H. armigera* larvae. (D) Binding of *HaTPS/TPP* promoter DNA with HaE2F transcription factor protein. Lane 1: *HaTPS/TPP* promoter DNA only (75 ng). Lane 2: HaE2F protein only 2 ug). Lane 3: DNA marker. Lane 4: *HaTPS/TPP* promoter DNA (75 ng) with HaE2F protein (0.5 ug). Lane 5: *HaTPS/TPP* promoter DNA (75 ng) with HaE2F protein (1 ug). Lane 6: *HaTPS/TPP* promoter DNA (75 ng) with HaE2F protein (2 ug). Lane 7: *HaTPS/TPP* promoter DNA (75 ng) with HaE2F protein (4 ug). Lane 8: *HaTPS/TPP* promoter DNA (75 ng) with HaE2F protein (8 ug). The gel shown in the upper panel was stained with SYBR Green EMSA stain. The gel shown in the lower panel is the same gel stained with SYPRO Ruby EMSA stain. Asterisks indicate the significance level between control and treatment. Student’s unpaired t-test was used for calculating significance. Data are represented as mean ±SEM; (*p<0.05; **p value <0.01; *** p value <0.001; ****p value <0.0001; ns=non-significant).

### 2.8 Dp silencing disrupts trehalose metabolism and myogenic gene regulation, affecting insect survival and eclosion

To study the regulation of trehalose metabolism and muscle development by E2F/Dp, we silenced the *Dp* (Zappia & Frolov, 2016) in *H. armigera* larvae. Silencing of *HaDp* resulted in a significant decrease in *HaE2F (∼*76%) and *HaDp* expression (∼56%) **(Figure 7A)**. The depletion of *HaE2F* and *HaDp* resulted in delayed pupation and pupal lethality **(Figure 7B)**, and the survived moths showed abnormal phenotypes such as abnormal wings in the adult moths **(Figure 7C)**, suggesting a role of E2F/Dp in eclosion. In insects, adult skeletal muscles are formed during the pupal stage (Zappia & Frolov, 2016), and trehalose and glucose are necessary for the growth and development of adult structures during this stage (Matsuda et al., 2015). Thus, we analyzed the expression level of trehalose metabolism genes, *TPS/TPP*, *Treh1*, and myogenic genes, *Prm* and *Tm2* upon silencing of *HaDp*. Interestingly, *HaTPS/TPP*, *HaPrm*, and *HaTm2* showed significantly reduced expression whereas *HaTreh 1* showed significant upregulation in *HaDp-*depleted insects **(Figure 7D)**, consistent with the previous report in *D. melanogaster* (Zappia et al., 2019, 2023b). The increase in the expression of *Treh* can be due to the utilization of available trehalose for survival in energy-depleted conditions (Shukla et al., 2015). These results suggest an essential role of *E2F/Dp* in metabolism and muscle development, which is crucial for animal viability and successful eclosion.

**Figure 7.**
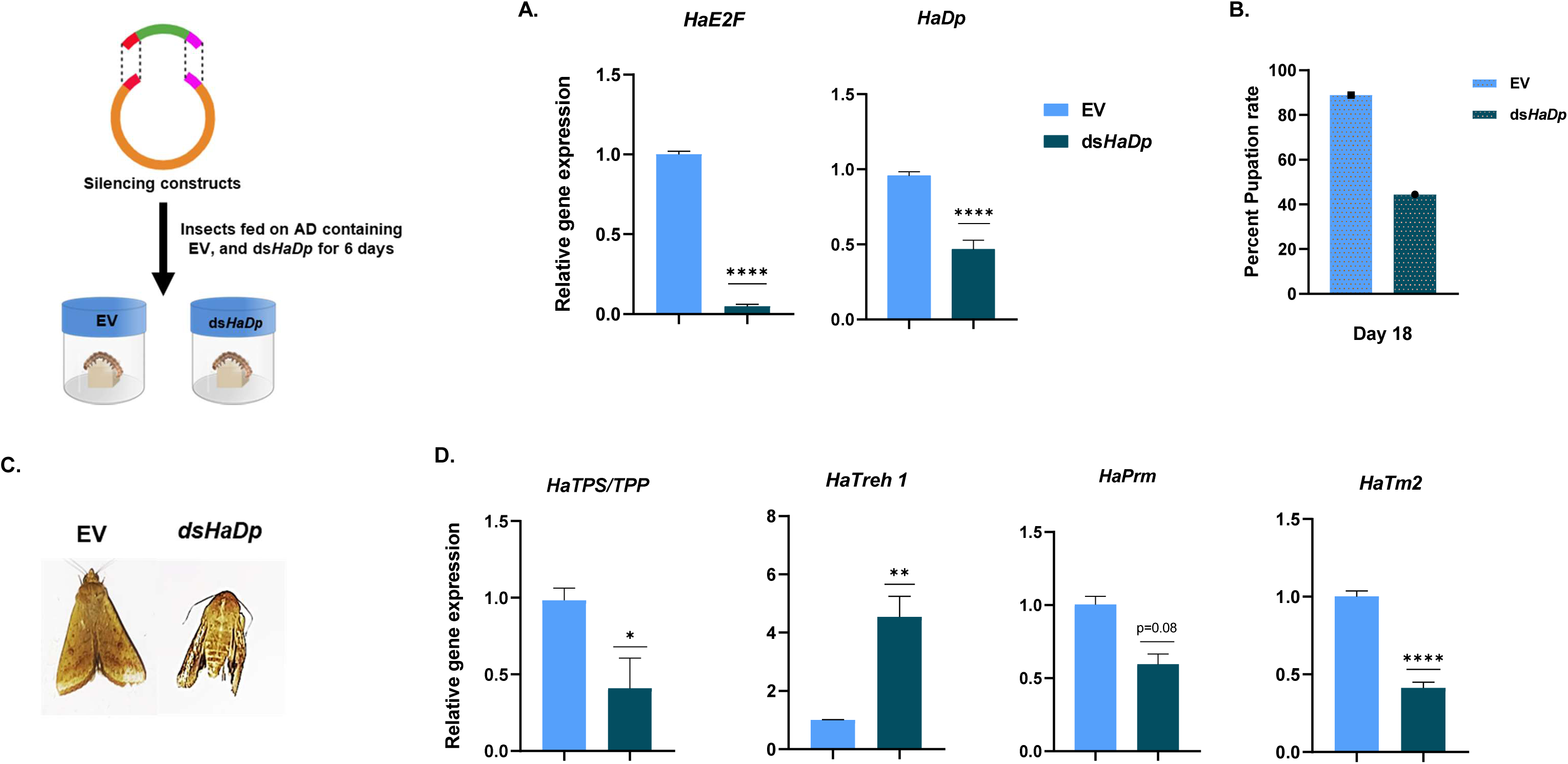
dsRNA mediated silencing of *H. armiger*a *HaDp* to study the role of E2F in metabolism and muscle development (n=36, Second instar larvae). (A) Silencing confirmation of *HaE2F and HaDp* using qRT-PCR. (B) Percent pupation rate calculated on the 18^th^ day of assay. (C) Phenotypic abnormalities associated with moths were observed upon *dsHaDp* feeding. (D) Gene expression analysis of trehalose metabolic genes and myogenic genes by qRT-PCR

### 2.9 Exogenous trehalose feeding mitigates the lethal effect of *HaE2F/*Dp-depletion in *H. armigera*

To investigate the regulation of *E2F/Dp* by trehalose, we examined whether exogenous trehalose supplementation could rescue the lethal effects of *HaDp* silencing on pupation and insect viability. Second instar larvae of *H. armigera* were initially fed with either an EV or *dsHaDp* to silence *HaE2F/Dp*. Subsequently, both EV and ds*HaDp*-fed insects were provided with 50 mM trehalose. Notably, exogenous trehalose feeding resulted in a significant increase in body weight of ds*HaDp*-fed insects while no change in the body weight observed in EV-fed insects **(Figure 8A-B)** and pupation rates **(Figure 8C)** in *HaDp*-silenced insects, along with the successful emergence of adult moths **(Figure 8D)**. Expression analysis revealed a significant upregulation of *HaE2F*, *HaDp*, and myogenic genes (*HaPrm* and *HaTm2*) and upregulation in trehalose metabolism genes (*HaTPS/TPP*, *HaTreh1*), in *HaE2F/Dp*-silenced insects upon exogenous trehalose feeding **(Figure 8E)**. Additionally, we assessed gene expression in insects fed an artificial diet (AD) and AD with 50 mM trehalose. Here, *HaTPS/TPP* expression was significantly downregulated, potentially due to feedback inhibition by exogenous trehalose, while *HaE2F*, *HaDp*, *HaTreh1*, *HaPrm*, and *HaTm2* were upregulated considerably **(Figure 8–figure supplement 5B)**. Despite the lack of a notable change in pupation rates **(Figure 8-figure supplement 5A)**, insects fed with trehalose exhibited normal eclosion **(Figure 8D)**. These findings demonstrate that trehalose levels regulate the expression of *E2F/Dp*, which is crucial for muscle development and successful eclosion.

**Figure 8.**
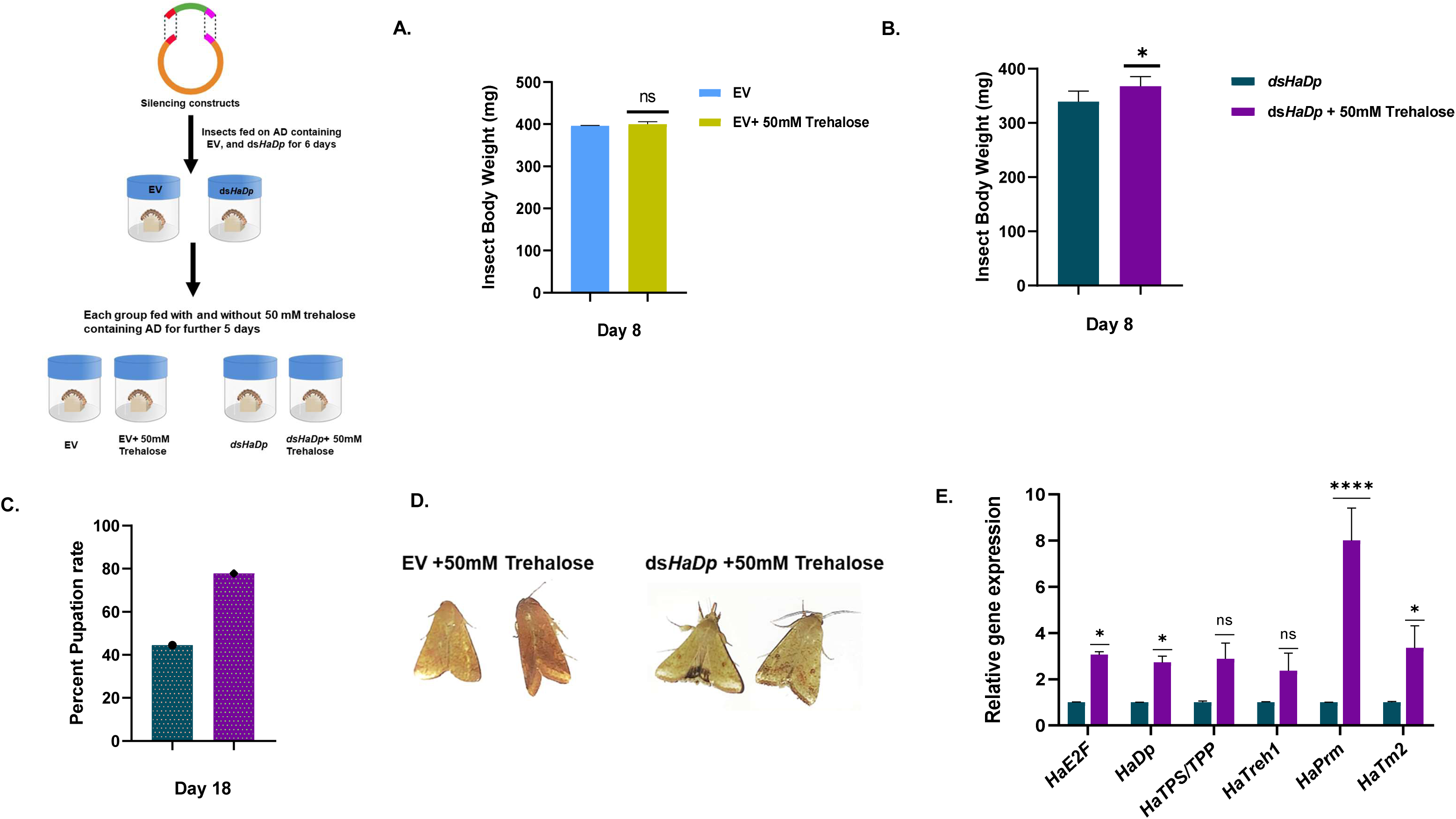
Exogenous feeding of 50 mM trehalose to the *HaDp*-silenced insects. Schematic representation of rescue assay (n= 36, second instar larvae). (A) Insect body weight upon exogenous feeding of 50 mM trehalose to EV-fed insects. (B) Insect body weight upon exogenous feeding of 50 mM trehalose to ds*HaDp*-fed insects. (C) Percent pupation rate calculated 18^th^ day of assay. (D) Normal adult moth emergence upon exogenous feeding of 50 mM trehalose to *HaDp*-silenced insects. (E) The gene expression level of TFs *(HaE2F*, *HaDp),* trehalose metabolic genes (*HaTPS/TPP, HaTreh1*), and myogenic genes (*HaPrm, HaTm2*) by real-time PCR upon exogenous feeding of 50 mM trehalose to *HaDp*-silenced insects. Asterisks indicate the significance level between control and treatment. Student’s unpaired t-test was used for calculating significance. Data are represented as mean ±SEM; (*p<0.05; **p value <0.01; *** p value <0.001; ****p value <0.0001; ns=non-significant).

### 2.10 Exogenous trehalose feeding partially alleviates the effects of trehalose synthesis inhibition in *H. armigera*

Since inhibiting trehalose synthesis affects the eclosion process by dysregulating TFs and genes involved in muscle development and function, we hypothesized that exogenous trehalose feeding could counteract the effects of trehalose synthesis inhibition. To test this hypothesis, we fed 50 mM trehalose to *HaTPS/TPP*-silenced insects exogenously and studied its effect on the expression level of TFs and myogenic genes responsible for muscle development and eclosion. Silencing *HaTPS/TPP* led to a significant reduction in *HaTPS/TPP* expression and activity **(Figure 9A)**, confirming the successful silencing of *HaTPS/TPP*. Although no significant changes in body weight or size were observed after feeding 50 mM trehalose to *HaTPS/TPP*-silenced insects **(Figure 9-figure supplement 6A-B)**, we noted a non-significant increase in *TPP* and *Treh1* expression and activity on the 10th day post-feeding **(Figure 9B)**. This suggests that exogenous trehalose may partially compensate for the silencing of *HaTPS/TPP*, thereby affecting trehalose metabolism. To further investigate, we analyzed the expression levels of transcription factors and myogenic genes. Interestingly, *HaE2F*, *HaDp*, *HaMef2*, and its target gene *HaPrm* were upregulated **(Figure 9C**). Although *HaMlx*, *HaMlxIP*, and *HaTm2* did not show significant changes in expression, the results still suggest a positive regulatory effect of exogenous trehalose on genes related to muscle development. Furthermore, we compared gene expression between insects fed with an artificial diet (AD) and AD with 50 mM trehalose. Here, *HaTPS/TPP* showed significant downregulation. While, *HaTreh1*, *HaE2F*, *HaDp*, *HaMef2*, *HaMlx*, *HaMlxIP*, *HaPrm*, and *HaTm2* were significantly upregulated **(Figure 9-figure supplement 6C)**. These findings support the hypothesis that exogenous trehalose feeding partially rescues the inhibition of trehalose synthesis and highlights its role in insect muscle development via transcriptional regulation.

**Figure 9.**
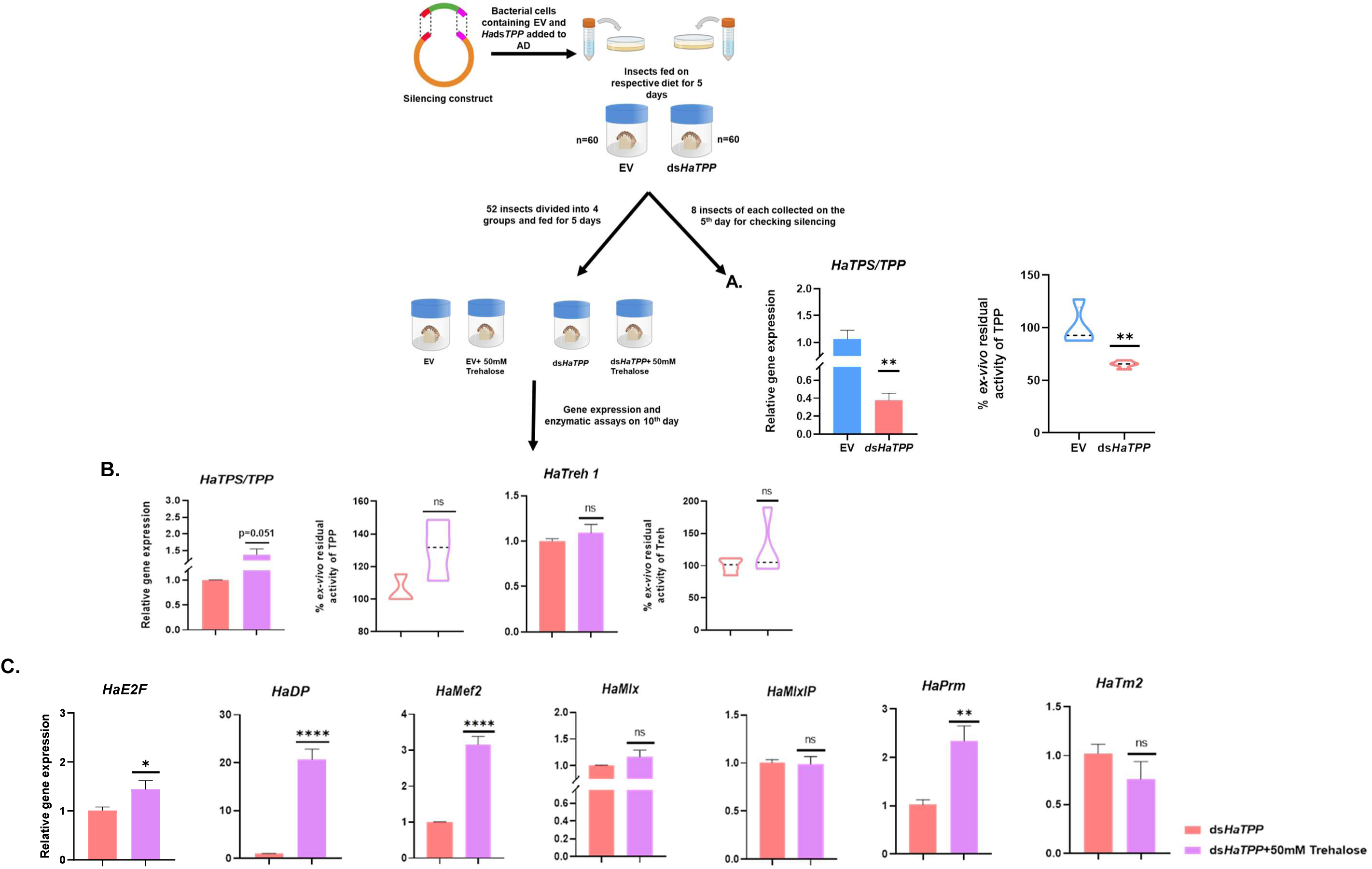
Exogenous feeding of 50 mM trehalose leads to partial rescue of the *HaTPS*/*TPP*-silenced insects. Schematic representation of rescue assay (n= 60, second instar larvae). (A) Silencing confirmation and *ex-vivo* residual activity of *Ha*TPP. (B) Gene expression analysis and residual activities of *Ha*TPP and *Ha*Treh compared between *dsHaTPP* and *dsHaTPP* plus 50 mM trehalose feeding. (C) Gene expression analysis of myogenic genes and selected TFs upon exogenous feeding of 50 mM trehalose to *HaTPS/TPP*-silenced insects. Asterisks indicate the significance level between control and treatment. Student’s unpaired t-test was used for calculating significance. Data are represented as mean ±SEM; (*p<0.05; **p value <0.01; *** p value <0.001; ****p value <0.0001; ns=non-significant).

## 3. Discussion

Trehalose serve as a rapid energy source during periods of high demand, such as metamorphosis and eclosion (Becker et al., 1996; Tang et al., 2018). While its role in regulating insect growth is well-established, its function in muscle development remains elusive. Disruption of trehalose synthesis, whether through silencing of *TPS/TPP* or by inhibition, results in broad energy depletion, activation of stress pathways, and collapse of myogenic programs (Bretscher & O’Connor, 2020; Matsuda et al., 2015; Sacktor & Wormser-Shavit, 1966; Tellis et al., 2024).

Differential gene expression analysis revealed that most myogenic genes showed reduced expression when trehalose synthesis was inhibited. Additionally, qRT-PCR results demonstrated significant overexpression of *HaTPS/TPP* in response to NPP treatment, likely due to feedback regulation. In contrast, *HaTreh1* and *HaTreh2* were significantly downregulated, possibly due to reduced trehalose availability. Moreover, various myogenic genes, including *HaPrm, HaTm2, HaFln,* and *HaMLC2* exhibited altered expression levels following *Ha*TPP inhibition. Furthermore, silencing of *TPS/TPP* leads to various adverse effects, including pupal defects, late pupal mortality, and unsuccessful eclosion. The fifth-instar larvae showed molting failures, leading to death, while rescued adults exhibited abnormal wing shapes and difficulties shedding their cuticles during the pupal stage. These findings are consistent with previous studies in *D. melanogaster* (Matsuda et al., 2015) and *Diaphorina citri* (Liu et al., 2020), further underscoring the critical role of trehalose metabolism in insect development. Histological analysis revealed fragmented muscle fibers in insects with inhibited trehalose synthesis, suggesting impaired muscle development. Silencing *HaPrm* led to outcomes similar to *dsHaTPP* fed insect, such as reduced survival, molting defects, deformed wings and fragmented muscles. These observations align with findings from studies on *Trichinella spiralis* paramyosin (*Ts-Pmy*), reinforcing the importance of *Prm* in larval muscle growth (X. Chen et al., 2012; Makovický et al., 2020). The impairment in muscle development was confirmed by the reduced expression of myogenic genes upon *TPS/TPP* and *Prm* silencing.

Moreover, dysregulation of trehalose metabolism led to the overall deregulation of genes involved in trehalose, glycolytic, and hexosamine pathway, resulting in reduced levels of glucose, glucosamine, glucosamine 6-phosphate, and energy molecules such as ADP, AMP, NAD, NADH and NMN. Loss of this flux triggers energy stress, reflected by elevated AMP levels and compensatory induction of arginine kinase, but these responses are insufficient to sustain normal growth. These results are consistent with previous reports from *H. armigera*, showing the importance of trehalose in maintaining energy homeostasis which is required for proper growth and development of insects (Tellis et al., 2024). We observed that trehalose-depleted insects exhibited significant reductions cell cycle related genes (e.g. SCAPER, *cyclin-dependent kinase 1*, *cyclin-A2-like*, and *G2/mitotic-specific cyclin-B, E2F/Dp)*, leading to deregulation of cell cycle. This indicates that the energy deficits might influence cell cycle progression and key cell proliferation mechanisms (Tixier et al., 2013; Zappia et al., 2019; Zappia & Frolov, 2016). Furthermore, we investigated glucose metabolism and muscle development related transcription factors such as *Mef2*, *Mlx* and its partner *MlxIP*, *IGF2*, and *Foxo* (Demontis & Perrimon, 2009; Hunt et al., 2015; Torrente et al., 2020). Following *TPS/TPP* silencing, these transcription factors exhibited altered expression, further supporting the link between trehalose synthesis and muscle development.

Further, GSEA data showed the upregulation of the processes linked to degradation of proteins (via proteasome) and alternative carbohydrate sources to get catabolized through energy metabolic processes thereby compensating for the defect in energy production and protein synthesis caused by *TPS/TPP* silencing. It could be possible that proteasome activity is linked to motor proteins as proteins targeted for degradation might be transported to proteasome through motor proteins. As the system is focused on recuperating with energy loss due to less availability of trehalose, a shift is observed from anabolism to catabolism. Thus, anabolic pathways like pentose-glucuronate conversions and retinol metabolism are downregulated.

At the transcriptional level, E2F/Dp emerges as a nodal regulator connecting trehalose metabolism to myogenic gene expression. Gene regulatory network inference and promoter-binding assays indicate that E2F/Dp binds not only muscle structural genes but also trehalose metabolism genes, positioning it as both a sensor and effector. The reduced expression of *E2F* and its dimerization partner, *Dp* upon inhibition of trehalose synthesis and silencing of *HaTPS/TPP* and *HaPrm* and also the presence of E2F binding sites on the promoters of *TPS/TPP, Treh1, Pgk, Act88F, Fln,* and *Prm,* suggested possible involvement of *E2F/Dp* in metabolism and muscle development. This conclusion is supported by the silencing of *E2F/Dp* in *H. armigera* led to late pupal mortality, and the emergence of adults with abnormal phenotypes, and reduced the expression of trehalose metabolism and myogenic genes. This suggests that E2F may regulate both trehalose metabolism and muscle development. EMSA supported direct binding of the E2F DNA-binding domain to the *HaTPS/TPP* promoter. Our findings are consistent with studies in *D. melanogaster*, where E2F regulates myogenic, glycolytic, and mitochondrial genes during pupal development (Zappia et al., 2019; Zappia & Frolov, 2016). ChIP data confirmed E2F binding on the promoters of glycolytic genes, such as *Pgk*, and myogenic genes, including *Mef2, Fln, Act88F*, and *Tm1* (Zappia & Frolov, 2016). The deficiency in E2F regulation on the *Pgk* gene was associated with increased mortality due to defects in energy-demanding tissues like muscles (Zappia et al., 2023a). Moreover, E2F depletion reduced muscle gene expression, emphasizing its role in muscle growth and development. The GSEA, GRN and gene expression analyses strongly suggest trehalose metabolism coordinates metabolic state with transcriptional programs that control developmental muscle architecture.

Silencing of *TPS/TPP* caused a reduction in trehalose and glucose levels, essential energy sources for muscle function and development. Interestingly, the exogenous feeding of trehalose to *E2F/Dp*-silenced insects showed increased pupation rate, and expression level *E2F* and *Dp*, trehalose metabolism genes and myogenic genes and successful emergence of adult moths, suggesting the regulation of *E2F/Dp* by trehalose. Similarly, exogenous feeding of trehalose to *TPS/TPP*-silenced *H. armigera* partially restored trehalose metabolism and the expression of *E2F*, *Dp*, and myogenic genes. The increased expression of *E2F*, *Dp*, *Mef2*, and *Prm* upon exogenous trehalose feeding further supports the hypothesis that E2F may regulate muscle gene expression in response to trehalose availability during eclosion. Despite partial rescue, the incomplete recovery could be due to the low absorption rate of exogenous trehalose through the gut. This supports a model in which trehalose metabolism, E2F/Dp activity, and sarcomeric assembly are interdependent processes that must be tightly coordinated for successful development. The integration of energetic supply with transcriptional regulation through this axis underscores how metabolic status is translated into tissue-specific developmental outcomes.

The most significant limitation is the unresolved causal link between metabolic flux and E2F regulation. Although reduced *E2F/Dp* expression was observed upon *TPS/TPP* silencing and partial rescue occurred with trehalose supplementation, the direct molecular signal connecting energy depletion to transcription factor repression remains unclear. Also, whether E2F/Dp senses metabolite levels directly or is modulated through upstream signaling pathways such as AMPK or TOR will require additional investigation. Dissecting signaling pathways that link energy depletion to E2F repression through genetic or pharmacological perturbation of AMPK and TOR could clarify whether these act as intermediaries. Furthermore, metabolomic analyses provided endpoint snapshots but did not capture dynamic fluxes that are central to establishing causality between trehalose hydrolysis and glycolytic throughput. In addition, while our network analysis highlighted E2F/Dp as central, other transcription factors responsive to glucose and energy status, including Mlx/MlxIP and Mef2, were also regulated. Their exact contribution and interplay with E2F/Dp remain to be mapped in detail. Despite these limitations, our work positions trehalose metabolism as more than a passive energy reservoir. By sustaining glycolytic flux and engaging E2F/Dp-dependent transcriptional control, trehalose metabolism ensures proper muscle development and successful eclosion. This study laid foundation to understand how insects adapt developmental timing to nutritional environments.

## 4. Material and Methods

### 4.1 Vector construction and dsRNA preparation

To carry out RNA interference (RNAi), specific partial fragments of *HaTPS/TPP* (490 bp), *HaPrm* (490 bp), *SfTPS/TPP* (490bp) and *HaDp* (490bp) were cloned into the L4440 vector between *EcoRI* sites using In-fusion cloning (Takara Bio). The resulting recombinant vectors, namely *dsHaTPP*, *dsHaPrm,* ds*SfTPP* and ds*HaDp* were confirmed by sequencing. Next, the prepared dsRNA vectors and L4440 (EV) were transformed into *E. coli HT115* (DE3) cells, which are deficient in RNAaseIII. A single colony from each transformation was selected and grown overnight in LB media containing ampicillin (100 ug/ml) and tetracycline (12.5 ug/ml) at 37 °C with shaking at 180 rpm. The overnight cultures were then used to inoculate fresh 200 ml 2X YT broth at a 1:100 proportion, along with the appropriate antibiotics. The cells were further grown at 37 °C until the optical density (OD) reached 0.5. At this point, the expression of double-stranded RNA (dsRNA) was induced by adding 1mM Isopropyl β-D-1-thiogalactopyranoside (IPTG), and the cells were allowed to continue incubating at 37 °C for 4-5 hours. To verify the successful expression of dsRNA, total RNA was extracted from induced and uninduced *E. coli HT115* cells using TRIzol reagent Invitrogen, USA), following the manufacturer’s instructions. The dsRNAs were then visualized using a 1.5% agarose gel. Finally, the cells were harvested by centrifugation at 5000g for 5 minutes at 4°C, and the pellet was resuspended in 5 ml of distilled water. This prepared the dsRNA constructs for further use in RNAi experiments (**Figure 2, 3-figure supplement 1**).

### 4.2 Feeding bioassay of NPP, *dsHaTPP, dsHaPrm*, ds*SfTPP,* and ds*HaDp*

In N-(phenylthio) phthalimide (NPP) inhibition bioassay, second instar larvae of *H. armigera* (n=100) were fed on an artificial diet (AD) containing 1000 ppm of NPP (TCI, Tokyo, Japan). Growth, development, body weight, and survival were analyzed and assessed, every alternate day for 10 days.

In the RNA interference (RNAi) experiments, second-instar larvae of *H. armigera* and *S. frugiperda* (n=60) were used for the feeding assay. The bacterial cultures containing dsRNA targeting *TPS/TPP* and L4440 empty vector (EV) were used for feeding *H. armigera* and *S. frugiperda* larvae while dsRNA targeting *HaPrm*, *HaDp* and EV were used to feed *H. armigera* larvae. The bacterial cultures were induced with IPTG, harvested by centrifugation, and resuspended in distilled water. The AD was prepared as per Chikate et al., 2016 (Chikate et al., 2016), and the resuspended bacterial cultures were added to the diet to achieve a final concentration of OD 1. A diet containing the EV served as the control, while the AD containing *dsHaTPP*, ds*SfTPP, dsHaPrm,* and ds*HaDp* fed to the insects on alternate days for a total of 6 days. Insect growth and survival on the control and dsRNA-containing diets were monitored every alternate day until pupation. On the 8th day, 30 insects from each treatment and control group were harvested and grouped in sets of ten to create three biological replicates. Additionally, 4 insects were fixed in 4% paraformaldehyde for microtomy to observe internal changes upon *dsHaTPP* and *dsHaPrm* feeding. The remaining insects were kept for further observations, including developmental abnormalities and eclosion. The silencing effects of *HaTPS/TPP, SfTPS/TPP, HaPrm* and *HaDp* were assessed by measuring larval mass gain, percent mortality, and the presence of phenotypic abnormalities, and analyzing the reduction of *TPS/TPP*, *Prm* and *E2F/Dp* transcript expression. The efficacy of *HaTPS/TPP, SfTPS/TPP, HaPrm,* and *E2F/Dp* silencing was determined using qRT-PCR analysis of selected genes.

### 4.3 Transcriptome analysis of insect tissue

RNA-Seq libraries were prepared for insects fed with EV, ds*HaTPP*, and ds*HaPrm* at the 8^th^ day post-feeding, with three biological replicates for ds*HaTPP* and two biological replicates for ds*HaPrm* treatment group (each replicate consisting of RNA pooled from ten insects). Total RNA was extracted using TRIzol reagent (Invitrogen, USA) following the manufacturer’s protocol. The RNA-seq libraries were constructed using the NEBNext Ultra II RNA Library Prep Kit for Illumina. These indexed libraries were then sequenced (2x 150bp) on a Novaseq 6000 platform at Lifecell International Pvt Ltd, Chennai. Raw RNA sequencing data is submitted to NCBI as a Bio project. (Bio Project IDs: PRJNA1176505 and PRJNA1337459).

### 4.4 RNA-sequencing data pre-processing

Raw FASTQ reads were aligned to the *H. armigera* reference genome (GCF_030705265.1) using STAR aligner v.2.7.11b (Dobin et al., 2013) for performing splice-aware alignment. Gene-level count matrices were generated using “– quantmode Gene Counts” in STAR. Genes with at least 10 read counts were retained for further analysis. The pipeline used for the methodology is provided in **Figure 5A**.

### 4.5 Differential gene expression and gene set enrichment analysis (GSEA)

Normalization and differential gene expression analysis was performed using DESeq2 v.1.48.2 implemented in R v.4.5.1 (Love et al., 2014). Genes were ranked based on the following metric sorted from positive to negative values: -

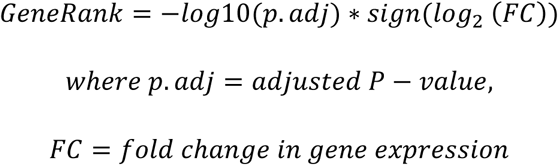

The ranked gene list was used to perform Gene Set Enrichment Analysis (GSEA) against the KEGG gene sets obtained from Enrichment Browser R package v.2.38.1 as the background. A minimum gene set size of 5 was considered for performing GSEA. Adjusted P-values were calculated using the Benjamini-Hochberg procedure. GSEA was performed using the fgsea procedure. GSEA was implemented using clusterProfiler v.4.16.0 in R. Gene sets with adjusted P-value < 0.05 were considered to be enriched.

### 4.6 Inference of the active gene regulatory network

Transcription factors in the *H. armigera* genome were searched using text-based Advanced Search from the NCBI Gene database using the query ‘(“Helicoverpa armigera“[Organism] OR “Helicoverpa armigera“[All Fields]) AND alive[prop] AND transcription factor’. The identified transcription factors (TFs) were further manually filtered to eliminate random matches with the text query. The shortlisted TFs and the normalized gene expression matrix were provided as an input to GENIE3 v. 1.30.0 for gene regulatory network inference, as implemented in R (Huynh-Thu et al., 2010; Su et al., 2010). The default parameters used for performing GENIE3 include the tree Method parameter which was set to Random Forest, number of candidate regulators randomly selected for best split determination (K) as square root and number of trees as 1000. The TF-target interactions were further filtered to retain only the top 10 percentile weighted interactions per target. This reduces the chance of false positives. We term the gene regulatory network obtained through this methodology as the “active gene regulatory network” (active GRN). The active GRN consists of source nodes which are TFs and the target nodes which are the targets that are predicted to be modulated by the TFs. The weighted edges represent the interaction of the TF node with the target node and the weight represents the strength of the interaction.

### 4.7 Functionally-enriched gene regulatory networks and network analyses

For obtaining gene regulatory networks of enriched processes, a subnetwork extraction approach was used. For subnetwork extraction, a list of all TFs along with genes belonging to each enriched gene set was prepared. Shortest path-based regulatory subnetwork extraction between the genes within this list from the active GRN was performed using the CytoHubba package v.0.1 in the Cytoscape Network Analyzer tool v.4.5.0. Shortest path biologically represents the transfer of information from TFs to targets via direct or indirect interactions. Genes appearing in this shortest path thus, are important in relaying regulatory information between functional nodes. Such subnetworks were extracted for every enriched KEGG gene set. We term these networks as “functionally-enriched GRNs” (fcGRNs) as these GRNs emerge upon TPS/TPP perturbation. Apart from individual gene sets, this analysis was repeated on the 11 gene sets together along with all the E2F family transcription factors from the text search-based TF list to generate an E2F transcription factor family - gene set subnetwork extraction, which we term as “E2F-fcGRN”. The fcGRN gives functional specificity and helps to find out TFs involved in every enriched function in an unbiased manner whereas the E2F-fcGRN provides information about how the E2F TFs are involved across all the gene sets combined. Network visualization and analysis was performed using Cytoscape v.3.10.2 (Shannon et al., 2003; Utriainen & Morris, 2023). The outdegree of every TF in the fcGRN and the combined-fcGRN was calculated using network analyses subroutine of Cytoscape.

### 4.8 Gene expression analysis

Differential gene expression analysis was performed for genes involved in trehalose metabolism, glycolysis, and muscle development from the *H. armigera* NPP, *dsHaTPP* and *dsHaPrm*-fed transcriptome. Heatmap was constructed in TB tools 1.09 (C. Chen et al., 2020). Clustering was performed using the Euclidean distance and complete clustering method. To assess the expression of selected genes, Real-time PCR (qRT-PCR) was performed following exposure to NPP at a concentration of 1000 ppm, dsRNA targeting *TPS/TPP* (*dsHaTPP, dsSfTPP*) and *Prm* (*dsHaPrm*) and *E2F/Dp* (ds*HaDp*). For analysis, three biological replicates of NPP treatment, ds*SfTPP* and *dsHaDp*, two biological replicates of *dsHaTPP*, and *dsHaPrm-*fed insects were used. Total RNA was extracted from insect tissue weighing between 80 to 100 mg using TRIzol reagent. To eliminate DNA contamination, the RNA was treated with an RQ1 RNase-free DNase enzyme. Subsequently, first-strand cDNA synthesis was performed using OligodTs primers following the manual of the High-Capacity cDNA Reverse Transcription Kit (Applied Biosystems, Foster, CA, USA), using 5 µg of RNA for a single reaction. The qRT-PCR analysis for transcriptomic validation was carried out using Takara TB Green Premix Ex Taq II (Tli RNaseH Plus, country) on a 7500 Fast Real-Time PCR System (Applied Biosystems, Foster, CA, USA). The primers used in the PCR reactions are listed in **Supplementary File Table 6**. Each PCR reaction was performed as three independent replicates. To determine the relative expression levels of the genes, the 2^_−_ΔΔCt method was applied, and *H. armigera Actin3ab* (*HaAct3ab*) and *S. frugiperda RPL10* (*SfRPL10*) were utilized as the endogenous reference gene to normalize the mRNA expression levels.

### 4.9 Residual enzyme activity of TPP and Treh

To extract protein from the whole insects of both control and treated larvae, the samples were crushed in liquid nitrogen. Approximately 150 mg of tissue was then suspended in 500 µl of 50 mM phosphate buffer at pH 7. The suspension was incubated at 4 °C for two hours, followed by centrifugation at 12,000 rpm for 20 minutes. The protein concentration was determined using the Bradford Reagent (Bio-Rad, Hercules, CA, USA) with bovine serum albumin (Sigma-Aldrich, MA, USA) as a standard. The enzymatic activity of TPP was assessed by measuring the release of Pi from trehalose 6-P using a malachite-green reagent (Sigma-Aldrich, MA, USA). The assay protocol was based on the method reported by Klutts et al., 2003, with some modifications. In a final volume of 100 µl, the reaction mixture contained 1 mM trehalose 6-P (Sigma-Aldrich, MA, USA), 2 mM MgCl_2_ (Hi-Media, MS, India), 50 mM sodium phosphate buffer at pH 7, and crude lysate protein (1 mg). The reaction mixture was incubated at 37 °C for 35 minutes. The reaction was terminated by adding two volumes of a filtered solution containing 0.15% malachite green, 1% ammonium molybdate (Hi-media, MS, India), and 12.5% concentrated HCl (Thomas Baker, MS, India) (Klutts et al., 2003) After allowing the mixture to develop color for 5-7 minutes at room temperature, the absorbance was measured at 630 nm.

For the trehalase activity assay, the release of glucose, a reducing sugar, from α, α-trehalose was measured using DNSA (Dinitro salicylic acid) reagent (Sigma-Aldrich, MA, USA). A total volume of 150 μL of trehalose at a concentration of 0.25% was added to a premix containing buffer and crude lysate protein (1mg). Following a 15-minute incubation, the reaction was stopped by adding 500 μL of DNSA reagent. The reaction tubes were then subjected to a boiling water bath for 5 minutes, and the absorbance was measured at 540 nm (Adhav et al., 2018).

### 4.10 Metabolomic analysis of NPP, *dsHaTPP,* and *dsHaPrm*-fed insects

To extract metabolites, the whole insect tissue from both control and treated samples was crushed, and approximately 80-100 mg of the tissue was used. Metabolite analysis was performed using a Q ExactiveTM Orbitrap mass spectrometer (ThermoFisher, USA) connected to a dual electrospray ionization (ESI) source. For metabolite separation, a Hypersil GOLD column (150 × 2.1 mm, 1.9 μm particle size) (Thermo Fisher, Vilnius, Lithuania) was employed in an ultra-performance liquid chromatography (UPLC) system. The separation was carried out at 35 °C with a flow rate of 0.2 ml/min. During the UPLC analysis, a 20-minute gradient was applied, with Solvent A consisting of 100% MS-grade water and Solvent B consisting of 100% MS-grade acetonitrile, both containing 0.1% formic acid. For data processing, MS-DIAL software (https://prime.psc.riken.jp/compms/msdial/main.html) was utilized. The software facilitated peak extraction, baseline filtering, calibration, peak alignment, deconvolution, peak identification, and integration. The LC-MS/MS spectral database of the Mass Bank of North America (https://mona.fiehnlab.ucdavis.edu/) was used for peak identification. The analysis focused on quantifying metabolites involved in trehalose metabolism, including UDP-glucose, glucose 6-phosphate, trehalose 6-phosphate, trehalose, and glucose.

### 4.11 Histological analysis of L4440 (EV), *dsHaTPP,* and *dsHaPrm*-fed insects

The treated fifth instar larvae were fixed in 4% paraformaldehyde for 24 h. The fixed samples were prepared according to the standard method as described by Lussier et al., 2023 (Lussier et al., 2023). The prepared samples were used for longitudinal sectioning (∼8 um) using the rotary microtome (Leica RM2126, Wetzlar, Germany), stained with hematoxylin-eosin, and observed by a light microscope using 10X magnification (Olympus, BX51, Tokyo, Japan). The skeletal muscle (SM) of treated larvae were observed and compared to the control.

### 4.12 Transcription factor binding site prediction

Nucleotide sequences ranging from −2200 to +200 relative to the start site of selected genes were extracted. Putative promoter regions and transcription start sites were predicted using the Berkeley *Drosophila* Genome Project’s Neural Network Promoter Prediction tool (https://www.fruitfly.org/seq_tools/promoter.html), allowing for the identification of complete promoter sequences and transcription start sites. Transcription factor binding sites were further analyzed using the JASPAR database (https://jaspar.elixir.no/) (Castro-Mondragon et al., 2022).

### 4.13 Electrophoretic mobility shift assay

The interaction between the *Ha*E2F transcription factor and the *HaTPS/TPP* promoter was confirmed by electrophoretic mobility shift assay (EMSA). The DNA-binding domain sequence of *H. armigera* E2F was cloned into the pET28a vector using *BamHI* and *HindIII* sites. The confirmed construct was expressed in *E. coli Shuffle T7* cells upon IPTG induction, and the recombinant E2F protein was purified via Ni-NTA affinity chromatography. Promoter regions of *HaTPS/TPP* were amplified from *H. armigera* genomic DNA using Phusion High-Fidelity DNA Polymerase and purified with the Wizard SV Gel and PCR-C Clean-Up System. Sequencing confirmed the promoter sequences. EMSA was performed using the ThermoFisher Scientific EMSA kit according to the manufacturer’s instructions. Briefly, 75 ng of promoter DNA was incubated with varying amounts of E2F protein (0.5 µg, 1 µg, 2 µg, 4 µg, and 8 µg) at room temperature for 3 hours. Samples were resolved on a 1% agarose gel, run at 35 V for 4 h at 4 °C. After electrophoresis, DNA was stained with SYBR Green EMSA gel stain for 30 min and visualized under UV transillumination at 300 nm. In the same gel, the protein bands were stained with SYPRO Ruby EMSA stain (Thermo Fisher Scientific) following a trichloroacetic acid (TCA) for 3 hours, and subsequently destained and imaged at 300 nm UV transillumination.

### 4.14 Exogenous feeding of 50 mM trehalose to *HaTPS/TPP* and *HaDp*-silenced insects of *H. armigera*

This feeding assay was done to study whether the feeding of trehalose to *HaTPS/TPP* and *HaDp*-silenced insects rescues the silencing effect of *TPS/TPP* and *E2F/Dp*. Initially, second instar larvae of *H. armigera* (n=60 for *dsHaTPP*, n=36 for *dsHaDp*) were fed on AD containing either EV and *dsHaTPP and dsHaDp* for the first 5 days. Then On the 6^th^ day, EV, *dsHaTPP,* and *dsHaDp*-fed insects were transferred on the AD containing EV, EV+50 mM trehalose, ds*HaTPP*, and ds*HaTPP*+50 mM trehalose *dsHaDp,* and *dsHaDp*+50 mM trehalose and allowed to feed for a further 5 days, and on the 10^th^ day, some insects were harvested for further experiments. A few insects in the *dsHaDp* feeding assay were kept for observation up to eclosion. The rescue effect of *HaTPS/TPP and HaE2F/Dp*-silencing was assessed by analyzing the gene expression level and enzyme activity data.

### 4.15 Statistical analysis

All experiments were performed in triplicate. Data were expressed as mean ± SE using GraphPad Prism v8.0 (GraphPad Software, San Diego, CA, USA). The Students unpaired t-test and One-way ANOVA was used to analyze the statistical significance between groups. Asterisks indicate significant changes compared to control (*P-value < 0.05; **P-value < 0.01; ***P-value < 0.001, ****P-value < 0.0001).

## Acknowledgment and funding

Sharada Mohite and Tanaji Devkate acknowledges the University Grant Commission for Fellowship. The authors thank Mrs. Santhakumari and Dr. Mahesh Kulkarni for providing the mass spectrometry facility. The authors thank Dr. Meenakshi Tellis, Dr. Ranjit Barbole and Mr. Vineetkumar Nair for their suggestions and technical help. This work was supported by the Council of Scientific and Industrial Research (CSIR), India [YSA000826].

## Author contributions

Sharada Mohite, Conceptualization, Data curation, Formal analysis, Validation, Investigation, Visualization, Methodology, Writing - original draft, Writing - review and editing; Tanaji Devkate, Data curation, Formal analysis, Investigation, Visualization; Prashant Kalaskar, Data curation, Formal analysis, Investigation, Visualization; Prashant Singh, Formal analysis, Validation, Investigation; Abhishek Subramanian, Data curation, Formal analysis, Validation, Investigation, Writing - review and editing; Rakesh Joshi, Conceptualization, Resources, Formal analysis, Supervision, Funding acquisition, Investigation, Visualization, Methodology, Writing-original draft, Project administration, Writing - review and editing

## Data availability

The RNA sequencing data are available in the Sequence Read Archive public repository with the SRA accession number (PRJNA1176505: SRR31091193 to SRR31091198) and (PRJNA1337459: SRR35724962 to SRR35724967).

## Conflict of interest declaration

The authors do not have any conflict of interest of any type.

## Supporting Files

### Supporting information 1

**Figure Supplement 1.** Agarose gel of *dsHaTPP*, *dsHaPrm,* and *dsSfTPP* dsRNA expression in *E. coli HT115*. Related to **Figures 2, 3, and S3**

**Figure Supplement 2.** dsRNA mediated silencing of *HaTPS/TPP*, and *HaPrm* (A) Insect body weight, images of larvae, and percent-controlled mortality of *H. armigera* larvae calculated on the 8^th^ day of post-feeding of *dsHaTPP.* (B) Insect body weight, images of larvae, and percent-controlled mortality of *H. armigera* larvae calculated on the 8^th^ day of post-feeding of *dsHaPrm*.

**Figure Supplement 3.** dsRNA mediated silencing of *S. frugiperda SfTPS/TPP* (A) Silencing confirmation of *SfTPS/TPP* using qRT-PCR and percent *ex-vivo Sf*TPP residual activity of ds*SfTPP*-fed insects compared to EV. (B) Gene expression of analysis *SfTreh1* using qRT-PCR and percent *ex-vivo Sf*Treh residual activity of ds*SfTPP*-fed insects compared to EV. (C) Insect body weight, and images of larvae of *S. frugiperda* larvae calculated on the 6^th^ day of post-feeding of *dsSfTPP.* (D) Gene expression of selected myogenic genes using qRT-PCR upon silencing of *SfTPS/TPP*. (E) Gene expression analysis of shortlisted TFs, *SfMlx,* and *SfMlxIP* upon silencing of *SfTPS/TPP*.

**Figure Supplement 4.** E2F/Dp directly regulates the synthesis of trehalose (A) Putative transcription factor binding sites in *HaTPS/TPP*, *HaTreh1*, *HaPgk*, *HaAct88F, HaFln, HaTm1,* and *HaPrm* were predicted in +200 and −2,000 regions from the transcription start site (TSS) using the JASPER database. E2F binding sites were found to be abundant in putative promoter regions. (B) Expression and purification of DNA binding domain of HaE2F protein checked on 12% SDS-PAGE. (C) Western blot analysis of purified DNA binding domain of HaE2F protein.

**Figure Supplement 5.** Exogenous feeding of 50mM trehalose to *HaDp-*silenced insects. (A) Percent pupation rate calculated on the 18^th^ day of assay (B) Gene expression analysis of TFs (*E2F, Dp*), trehalose metabolic genes (*HaTPS/TPP*, *HaTreh1*), and myogenic genes (*HaPrm, HaTm2*) upon feeding AD-containing with EV, and EV+ 50mM trehalose.

**Figure Supplement 6.** Exogenous feeding of 50mM trehalose to *HaTPS/TPP*-silenced insects. (A) Insect body weight up to 6 days upon feeding AD-containing with EV, EV+ 50mM trehalose, *dsHaTPP*, and *dsHaTPP* +50 mM trehalose. (B) Larval images were taken on the 6^th^ day of assay. (C) Gene expression analysis of trehalose metabolism genes and selected TFs and myogenic genes compared between EV and EV+50 mM trehalose-fed insects.

### Supporting information 2

**Table S1.** Different gene expression analysis results from DESeq2

**Table S2.** Gene Set Enrichment Analysis with genes ranked according to the metric (GeneRank=-log10(p.adj)*sign(〖log〗_2 (FC)))

**Table S3.** Functional GRN edge lists for each enriched KEGG gene set with GENIE3 weights

**Table S4.** Analysis of Motor protein fcGRN

**Table S5.** The E2F family fcGRN

**Table S6.** List of primers used for gene expression analysis and cloning.

## Notes

### Competing Interest Statement

The authors have declared no competing interest.

